# Regenerative Signatures in Bronchioalveolar Lavage of Acute Respiratory Distress Syndrome

**DOI:** 10.1101/2023.11.13.566908

**Authors:** Runzhen Zhao, Marco Hadisurya, Harrison Ndetan, Nan Miles Xi, Sitaramaraju Adduri, Nagarjun Venkata Konduru, Buka Samten, W. Andy Tao, Karan P Singh, Hong-Long Ji

## Abstract

**Background:** In patients with severe acute respiratory distress syndrome (ARDS) associated with sepsis, lung recovery is considerably delayed, and mortality is much high. More insight into the process of lung regeneration in ARDS patients is needed. Exosomes are important cargos for intercellular communication by serving as autocrine and/or paracrine. Cutting-edge exomics (exosomal proteomics) makes it possible to study the mechanisms of re-alveolarization in ARDS lungs.

**Aims:** This study aimed to identify potential regenerative niches by characterizing differentially expressed proteins in the exosomes of bronchioalveolar lavage (BAL) in ARDS patients.

**Methods:** We purified exosomes from BAL samples collected from ARDS patients by NIH-supported ALTA and SPIROMICS trials. The abundance of exosomal proteins/peptides was quantified using liquid chromatography-mass spectrometry (LC-MS). Differentially expressed exosomal proteins between healthy controls and ARDS patients were profiled for functional annotations, cell origins, signaling pathways, networks, and clinical correlations.

**Results:** Our results show that more exosomal proteins were identified in the lungs of late-stage ARDS patients. Immune cells and lung epithelial stem cells were major contributors to BAL exosomes in addition to those from other organs. We enriched a wide range of functions, stem cell signals, growth factors, and immune niches in both mild and severe patients. The differentially expressed proteins that we identified were associated with key clinical variables. The severity-associated differences in protein-protein interaction, RNA crosstalk, and epigenetic network were observed between mild and severe groups. Moreover, alveolar type 2 epithelial cells could serve as both exosome donors and recipients via autocrine and paracrine mechanisms.

**Conclusions:** This study identifies novel exosomal proteins associated with diverse functions, signaling pathways, and cell origins in ARDS lavage samples. These differentiated proteins may serve as regenerative niches for re-alveolarization in injured lungs.

## Introduction

Sepsis is a leading cause of the acute respiratory distress syndrome (ARDS), contributing to approximately 45% of overall in-hospital mortality[1-3]. Similarly, the mortality rate of ARDS-associated septic shock and multiorgan failure approaches 30-50%[3-7]. ARDS is the most commonly caused by pneumonia (40%), including both viral and bacterial infections[1, 8], e.g., COVID-19[9]. Stem cell-based therapy holds the most substantial promise for the therapy of ARDS[10-16]. However, the role of the microenvironment (niches) in stem cell-mediated repair of injured lungs remains unknown.

Extracellular vesicles, including exosomes released from stem/progenitor cells, may serve as autocrine/paracrine mediators for repair. Exosomes are extracellular vesicles that originate from endosomes and contain miRNA, DNA, proteins, RNA, and subcellular compartments[17-19]. Accumulating evidence suggests that exosomes released from the stem and progenitor cells hold promise for the therapies of ARDS[20-30]. Exosomal “cytosolic” and surface membrane proteins surrounded by cellular lipids are directly derived from endosomes. The structure and function of exosomal proteins protected by lipid bilayers are least altered enzymatically compared with other lipid-free counterparts. To date, how exosomal signals in bronchioalveolar lavage (BAL) regulate the fate of stem cells during the lung regeneration remains obscure[31].

The predominant stem cells for re-alveologenesis include alveolar type 2 epithelial cells (AT2), bronchoalveolar stem cells (BASC), basal cells, endothelial progenitors, and mesenchymal stem cells (fibroblasts). These stem cells renew very slowly (> 6 months) to maintain their homeostasis in normal lungs. However, in injured lungs, they are activated to regenerate alveoli, and their fate is determined by the niches, including exosomal proteins[32, 33]. The Notch, Hedgehog, Wnt[12, 13], BMP, FGF, EGF, HGF, PECAM1, and histone deacetylase signaling pathways govern the fate and stemness of multipotent lung stem cells in vitro and in animal models[13, 34-38]. In addition, some stem cells in terminal airways (goblets, ciliated, and club cells) as well as inflammatory cells may contribute to alveolar repair. An emerging concept is that cytokines/chemokines could serve as niches for lung injury repair[39-41]. However, the stage specificity of the exosomal signals and their potential to act as biomarkers for ARDS patients have not been reported. The tendency to define stem cells by function is emerging[42, 43], but local stem cells and exosomal niches for repair at the late stage of septic lungs have not been completely analyzed for septic ARDS patients.

This study aimed to identify exosomal niches by analyzing BAL samples from ARDS patients. Our results demonstrate that these exosomal niches are specific to the condition’s severity. We observed differences between mild and severe patients in terms of the cell origins of BAL exosomes, enriched exosomal protein functions, and network interactions. This study also identified differentially expressed exosomal protein signatures correlated with numerous clinical variables.

## Methods

### Collection of clinical variables and BAL samples

All of the lavage samples and clinical data were provided by the BioLINCC, which were previously collected by NIH-supported multicenter, randomized, double-blind, placebo-controlled clinical trials (ALTA and SPRIOMICS)[44-47]. The institutional review board (IRB) of the University of Texas at Tyler Health Science Center (EXAMPT09-017) and Loyola University Chicago (LU#216964) approved the use of discarded BAL samples and deidentified clinical data. Patients were grouped based on the Berlin’s criteria for the severity of ARDS. The ratio of arterial oxygen tension (PaO_2_ in mmHg) to the fraction of inspired oxygen (FiO_2_) for each group is: PaO_2_/FiO_2_ 2300 for healthy controls, 200≤ PaO_2_/FiO_2_ <300 for mild, 100≤ PaO_2_/FiO_2_ ≤200 for moderate, and PaO_2_/FiO_2_ <100 for severe patients. The mild group is defined as the early stage of ARDS, often termed Acute Lung Injury (ALI). Standard NIH-recommended procedures were used for BAL collection, storage, and shipping. To avoid treatment-related biases, we included samples collected from placebo groups or those collected before treatments when available. Adjustments were made for treatment arms when pre-treatment samples were not available. Therefore, this study is a post-clinical trial project *per se*. The inclusion criteria for collecting BAL samples include: 1) the samples must have complete clinical records throughout the entire trials; 2) healthy controls must not have lung diseases and have a p/f ratio > 300 mmHg; 3) the BAL samples must have been collected from the placebo group or from the time points prior to the treatments; 4) the total volume of BAL samples must be > 0.5 ml; and 5) samples from all three stages of ARDS, as defined by the Berlin criteria must be available from the same clinical trial.

### Preparation of exosomal proteins

Exosomes from bronchioalveolar lavage (BAL) samples were captured and processed by Tymora Analytical Operations (West Lafayette, IN) using magnetic EVtrap beads as previously described[48]. The isolated exosomes were lysed to extract proteins using the phase-transfer surfactant (PTS)-aided procedure[48]. The proteins were reduced and alkylated by incubating them in 10 mM TCEP and 40 mM CAA for 10 minutes at 95°C. The samples were diluted five-fold with 50 mM triethylammonium bicarbonate and digested with Lys-C (Wako) at a 1:100 (wt/wt) enzyme-to-protein ratio for 3 hours at 37°C. Trypsin was then added at a final 1:50 (wt/wt) enzyme-to-protein ratio for overnight digestion at 37°C. To remove the PTS surfactants from the samples, trifluoroacetic acid (TFA) was added to the samples to a final concentration of 1% for acidification, and then mixed with an ethyl acetate solution at a 1:1 ratio. The mixture was vortexed for 2 minutes and then centrifuged at 16,000 × g for 2 minutes to separate the aqueous and organic phases. The organic phase (top layer) was removed, and the aqueous phase was collected. This step was repeated once more. The samples were then dried in a vacuum centrifuge and desalted using Top-Tip C18 tips (Glygen) following the manufacturer’s instructions. After desalting, the samples were completely dried in a vacuum centrifuge and stored at -80°C.

### LC-MS/MS analysis

Dried peptide samples were dissolved in 4.8 μL of 0.25% formic acid with 3% (vol/vol) acetonitrile and 4 μL of each sample was injected into an EasynLC 1000 (Thermo Fisher Scientific). Peptides were separated on a 45 cm in-house packed column (360 μm OD×75 μm ID) containing C18 resin (2.2 μm, 100 Å; Michrom Bioresources). The mobile phase buffer consisted of 0.1% formic acid in ultrapure water (buffer A), with an eluting buffer of 0.1% formic acid in 80% (vol/vol) acetonitrile (buffer B). The separation was achieved using a linear 60-minute gradient of 6–30% buffer B at a flow rate of 250 nL/min. The Easy-nLC 1000 was coupled online with a hybrid high-resolution LTQ-Orbitrap Velos Pro mass spectrometer (Thermo Fisher Scientific). The mass spectrometer was operated in the data-dependent mode, starting with a full-scan MS (from m/z 300 to 1,500 with a resolution of 30,000 at m/z 400), followed by MS/MS of the 10 most intense ions.

### LC-MS data processing

To quantify proteomics data, we extracted peptide intensities with the following parameters: initial precursor mass tolerance set at 10 ppm; minimum number of isotope peaks as 2; maximum ΔRT of isotope pattern multiplets at 0.2 minutes; PSM confidence FDR of 0.01; hypothesis test of ANOVA; maximum RT shift of 5 minutes; pairwise ratio-based ratio calculation; and maximum allowed fold change set at 100. To calculate fold changes between groups of the cohort, we summed total protein abundance values and compared the ratios of these sums across different samples. All protein and peptide identifications were grouped, and any redundant entries were removed. Only unique peptides and unique master proteins were reported.

### Identification of differentially expressed proteins in exosomes

The datasets generated from Proteome Discoverer were subjected to statistical analysis using Perseus, as described previously[49]. Protein abundance values were converted to log_2_ fold changes (FC), that is, the disease/control ratios (either averaged or matched). Proteins with at least 3 valid values per group were retained. Missing values were generated because the intensity of the proteins was too low to be detected by the instrument. Missing values were then imputed with small random values drawn from a normal distribution generated by Perseus with a downshift of 1.8 SDs and a width of 0.3 SDs. Data normalization was performed using the width adjustment method. A two-test sample t-test was performed to assess between-group differences. The resulting p-values and log_2_FC values were used to create volcano plots using the R EnhancedVolcano package. Normalized data was used to generate a heatmap using the R heatmap package. To minimize both false positive and false negative results, we applied the Benjamin and Hochberg method for q-tests to control the FDR. Both p- and q-values were evaluated across the entire list of values, rather than individually, to minimize the occurrence of false positives and negatives.

### Functional enrichment of differentially expressed proteins (DEPs)

We employed the EnrichR[50], ToppGene[51], and the LGEA Web Portal[52] were applied to track the cell origins of DEPs. Human was selected as the species, and all other default settings were used for functional annotations. GlueGO, a plug-in app for Cytoscape was utilized to visualize the network of DEPs[53, 54].

### Correlations of clinical variables and differentially expressed proteins

SAS (SAS Institute Inc.) was used to compute the correlations between exosomal proteins and clinical variables. Proteins with more than 2 missing values were discarded. The statistical association was noted when the p-value < 0.05. A multiple linear regression model was built to generate estimates for the clinical parameters and their corresponding 95% confidence intervals by adjusting for age, gender, fluid management, and baseline severity of illness.

## Results

### Baseline characteristics and clinical data of ARDS patients

The baseline of the cohort has been summarized in **Table 1**. A total of 89 clinical measures, including demographic information and vital organ function for all patients, were available for the samples. Detailed blood gas, tidal volume, minute ventilation, and mortality indices were quantitatively documented for up to 90 days. As shown in **Figure 1**, the BAL samples were processed to obtain pure exosomes, which were subjected to protein purification. Digested peptides were analyzed by LC-MS to generate proteomics datasets. These datasets were used to identify DEPs, track the cell origins of DEPs, perform functional enrichments, compute clinical correlations, and conduct network analyses.

**Figure 1.**
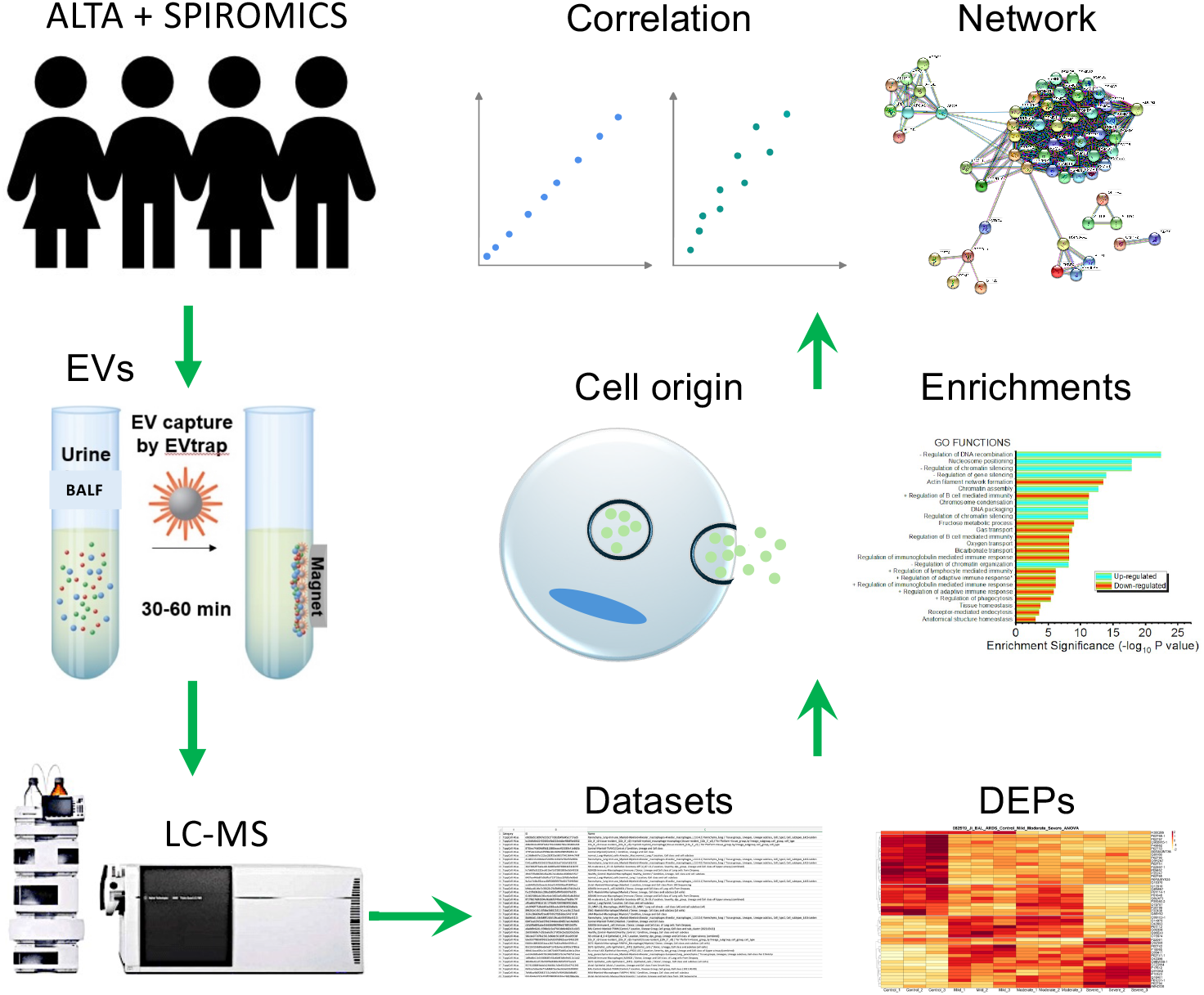
Workflow for the proteomics analysis of BAL exosomes. EVs, extracellular vesicles. DEPs, differentially expressed proteins.

**Table 1.** Demographics of control and ARDS cohorts. SD, standard deviation. GI, gastrointestinal.

|  | **Control** | **ARDS** |
| --- | --- | --- |
| Number of patients | 3 | 9 |
| Male | 2 | 5 |
| Female | 1 | 4 |
| Age (y ± SD) | 56.67 ± 4.93 | 40.00 ± 15.00 |
| White | 2 | 6 |
| Black | 1 | 2 |
| Hispanic | 2 | 1 |
| **Etiology** |  |  |
| Sepsis |  | 6 |
| Pneumonia |  | 6 |
| Aspiration |  | 4 |
| Trauma |  | 1 |
| Transfusions |  | 0 |
| **Sepsis site** |  |  |
| GI tract |  | 1 |
| Urinary tract |  | 1 |
| Unknown |  | 4 |
| **Mortality** |  |  |
| 60 days |  | 1 |
| 90 days |  | 1 |

### Differentially expressed proteins in BAL exosomes of ARDS

We have analyzed lavage samples from both healthy controls and ARDS cases (**Figure 2**). DEPs were compared between controls and the patients at different stages of the disease using heatmaps and volcano plots (**Figure 2A-F)**. In total, 4,458 exosomal proteins and 40,096 peptides were detected (**Figure 2G**). We observed both upregulated and downregulated exosomal proteins. More exosomal proteins were significantly altered in severe patients (612 DEPs) than those in the mild (386 DEPs) and moderate groups (334 DEPs). In addition, the ratio of differentially downregulated and upregulated proteins increased in a severity-dependent manner (1.05, 1.21, and 1.40, respectively, for mild, moderate, and severe patients). All three stages of ARDS shared considerable DEPs, either downregulated or upregulated (**Figure 2H & 2I**). These results suggest that a potential imbalance of downregulated and upregulated exosomal proteins may be associated with the severity of ARDS, supported by the evidence that the more severity the lung injury, the more exosomal proteins are downregulated.

**Figure 2.**
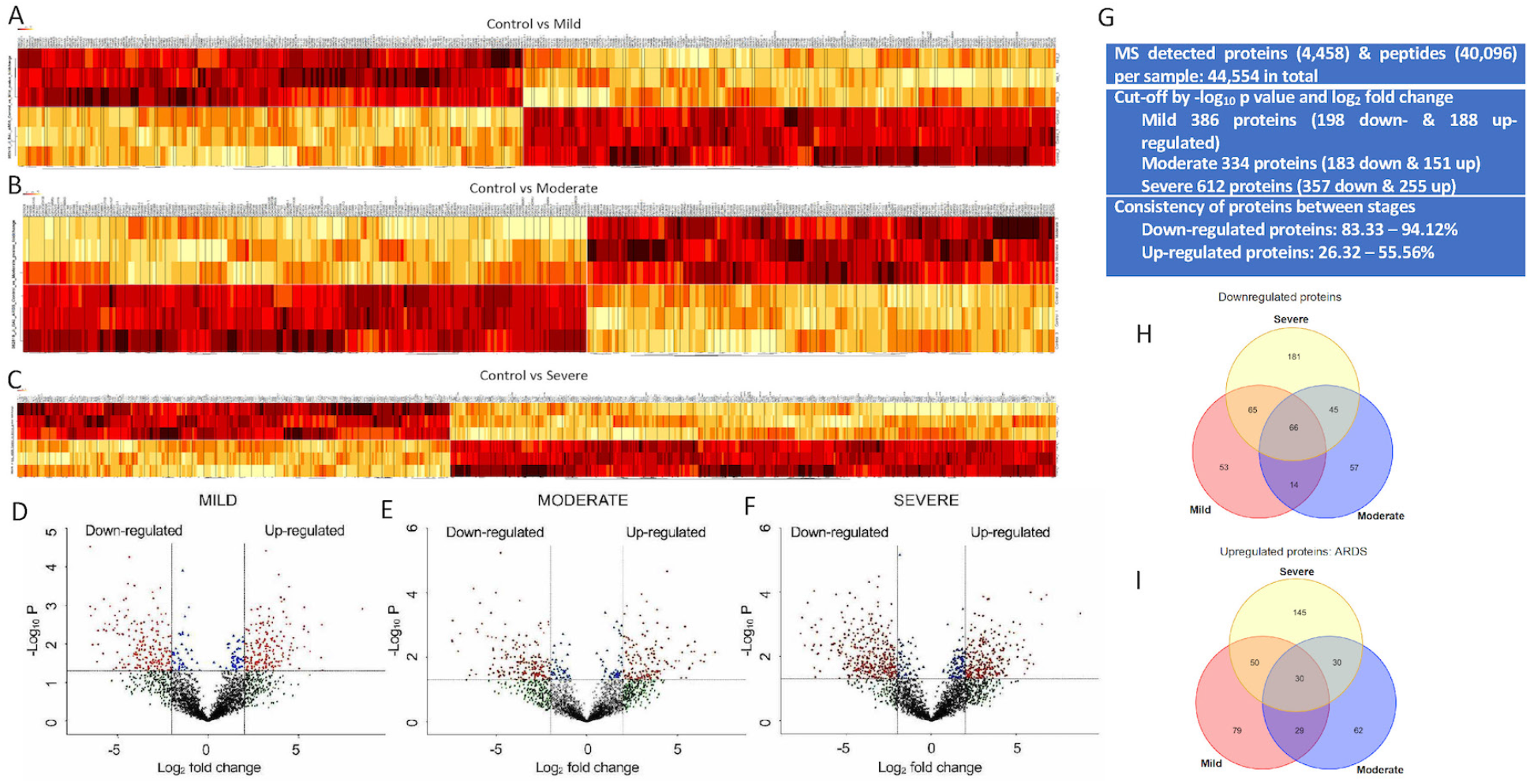
Identification of differentially expressed proteins. *A-C*. Heatmaps comparing upregulated and downregulated proteins between controls and three stages of ARDS. *D-F*. Volcano plots for the identification of DEPs for mild, moderate, and severe patients. Two cut-off criteria are: p-value < 0.05 and fold change > 2. *G*. Exosomal proteins identified by LC-MS and their consistency between stages. *H-I*. Venn plots showing the overlapping of downregulated (*H*) and upregulated (*I*) exosomal proteins. n = 12 patients.

### Cell origins of differentially expressed exosomal proteins

The blood-gas barrier is damaged in severe ARDS, allowing exosomes released from other organs to permeate into the airspaces via the circulatory system. Analysis of cell origin by the LGEA platform showed that the BAL exosomes were released from mesenchymal, immune, epithelial, and endothelial cells (**Figure 3A**). Epithelial cells accounted for approximately 50% of downregulated exosomal proteins in the severe group. In comparison, slight changes were observed in mesenchymal and endothelial cells for both downregulated and upregulated exosomal proteins. Furthermore, more specific cell subpopulations were identified using the ToppGene (**Figure 3B**). Severe patients had more cell populations than those in the mild group. Exosomal proteins from basal and epithelial cells were markedly reduced in severe patients. Additionally, in severe ARDS patients, additional exosomes were detected from the liver, cardiovascular system, bone marrow, pancreas, neurons, mesenchyme, and immune cells. Taken together, the top 5-ranked cell origins were macrophages, epithelial cells, myeloid cells, monocytes, and secretory cells (**Figure 3C**). Moreover, lung epithelial stem cells, including basal cells, club cells, and AT2 cells, significantly contributed to the DEPs in BAL exosomes.

**Figure 3.**
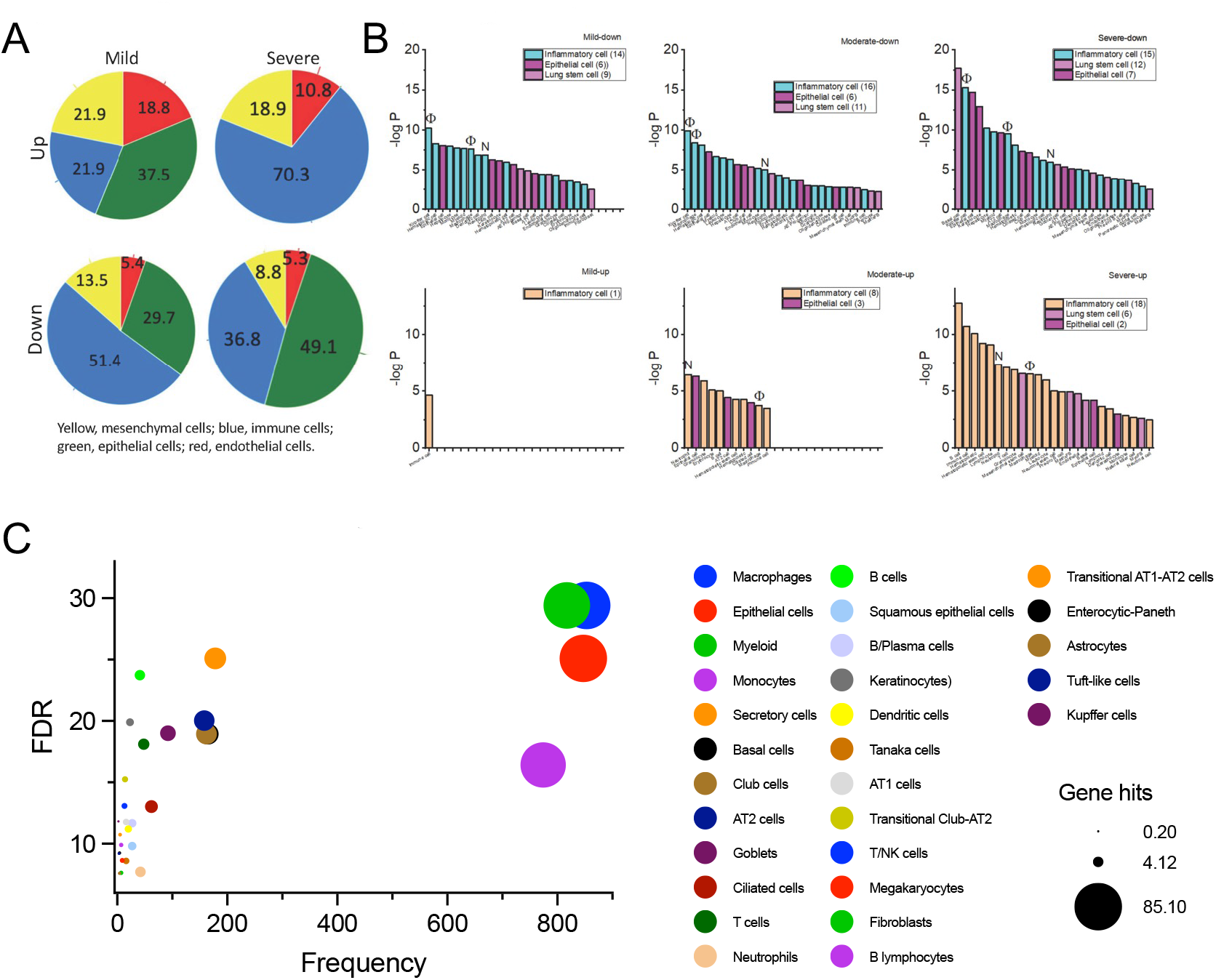
Cell origin of differentiated exosomal proteins. *A*. Analysis of the LGEA portal for significantly upregulated (Up) and downregulated (Down) exosomal proteins in both mild and severe patients. The percentage of contribution was shown for the origin of mesenchymal (yellow), immune (blue), epithelial (green), and endothelial (red) cells. *B*. Results from the ToppGene analysis. P-values of inflammatory cells, epithelial cells, lung stem cells were plotted against each cell sub-population. Total cell sub-populations for each category were given in brackets. N, neutrophil; Φ, macrophage. *C*. Summary of cell origin as a function of false detection rate (FDR).

### Diverse gene ontology between mild and severe ARDS

To examine the gene ontology (GO) of both downregulated and upregulated exosomal proteins in lavage samples, we performed the GO enrichment analysis. The top-ranked GO Biological Processes were shown in **Figure 4A**. A divergence was observed between mild and severe ARDS patients in the four top-ranked biological processes: 1) regulation of the biological process, 2) response to stimuli, 3) transport, and 4) cell organization and biogenesis. Significantly more exosomal proteins were downregulated than upregulated in severe ARDS patients. Neither upregulated nor downregulated biological processes differed significantly between the mild and moderate patients. Among the remaining functions, cell homeostasis that is balanced by cell lineage and death, predominated. Notably, coagulation plays a major role in exosomal proteins. Furthermore, GO Cellular Components, Molecular Function, and KEGG Signaling Pathway were visualized for exosomal DEPs. As shown in **Figure 4B**, five signaling pathways (hemostasis, complement cascade, innate immune system, phagosome, and neutrophil degranulation) and 25 interactive networks were enriched by the ClueGO for downregulated exosomal DEPs. Moreover, a pronounced difference in the GO terms was observed between downregulated (**Figure 4C**) and upregulated proteins (**Figure 4D**). These results demonstrate diverse molecular functions and signaling pathways between the early and late stages of ARDS patients.

**Figure 4.**
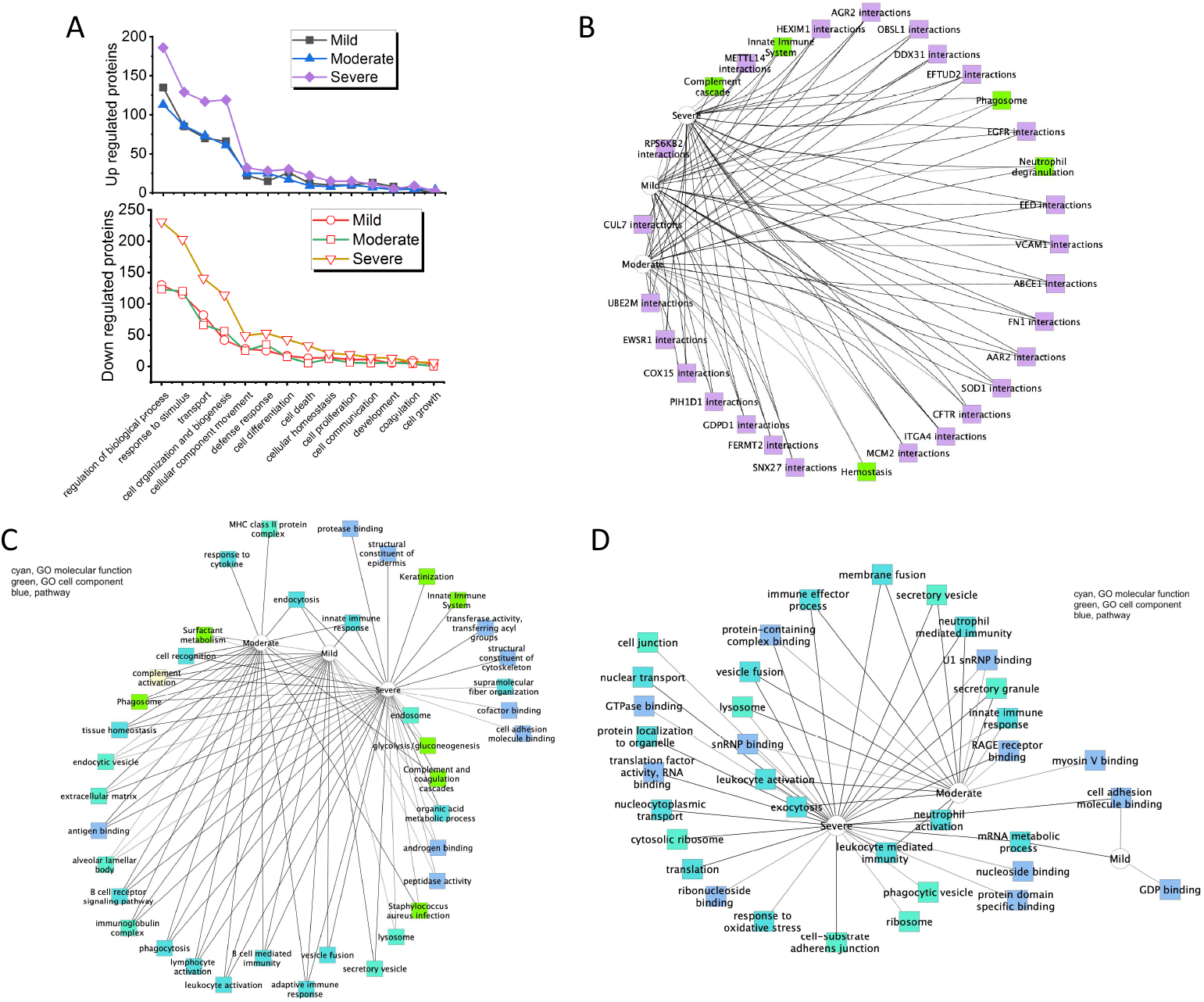
Enrichments of biological processes, functions, cell components, and signaling pathways of gene ontology (GO). *A*. Top 14 biological processes per ARDS stage for both upregulated (upper) and downregulated DEPs (bottom). *B*. Ontology of all downregulated proteins for mild, moderate, and severe groups. *C*. Ontology of all upregulated proteins for three groups. *D*. Ontology of common DEPs between all three stages of the disease.

### Dynamic signaling pathways for the regeneration of injured lungs

Based on the cell origin of BAL exosomal proteins, the lineages of stem cells could be regulated by traditional signaling pathways in AT2 cells, various growth factors, and immune niches. Considering the diverse GO enrichments between the early and later stages of ARDS patients (**Figure 4**), we compared the expression of differentiated exosomal proteins between the mild and severe groups. The SCF, FGFR, and PDGF signaling pathways were upregulated in mild patients (**Figure 5**). In contrast, the Notch3 cascade was enhanced in the severe group. The Hippo and Wnt signals were suppressed in mild patients. These severity-specific signal pathways may dynamically regulate lung stem cell-based re-epithelialization.

**Figure 5.**
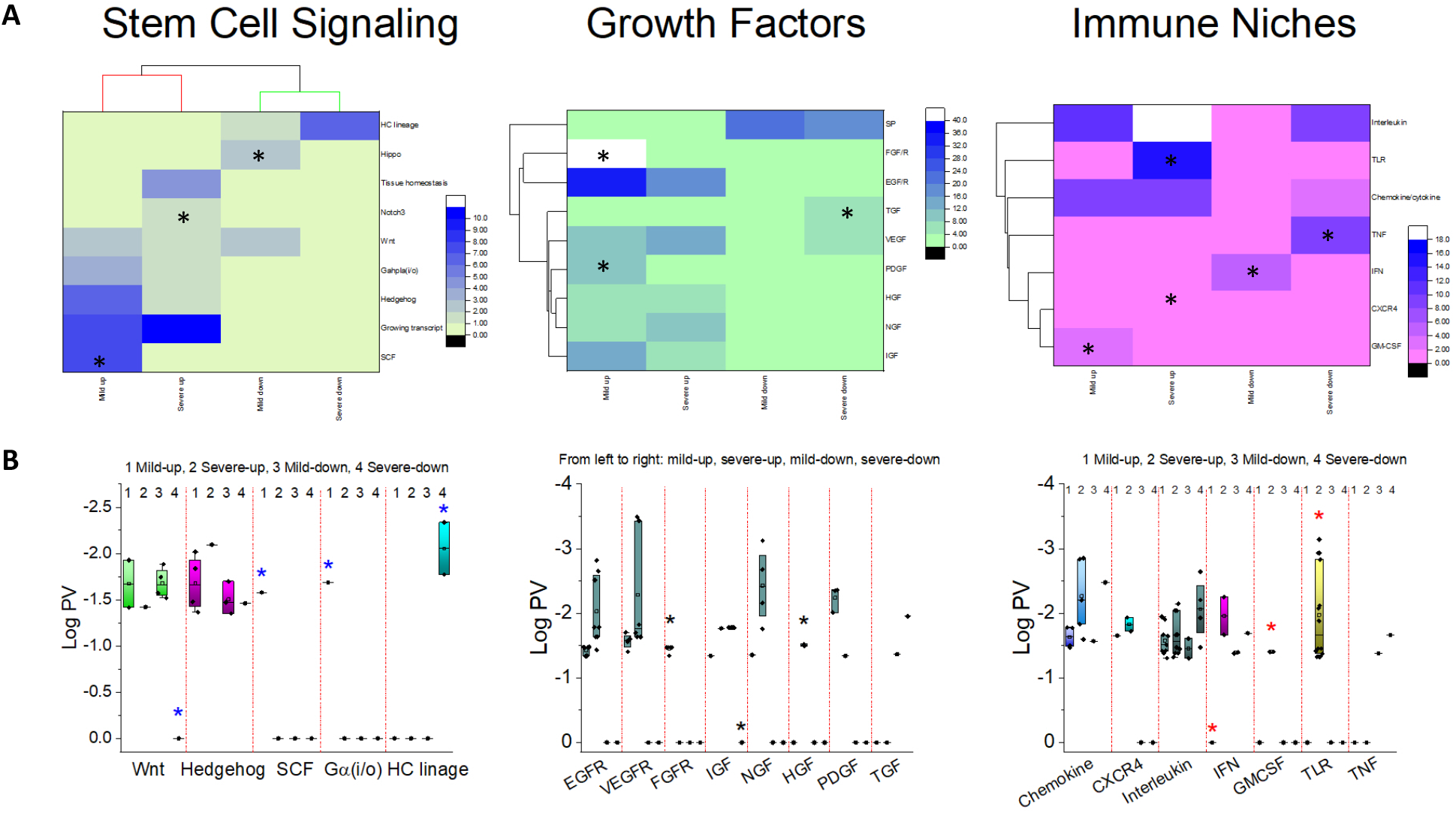
Stage-specific stem cell niches between early and later stages. *A*. Heatmaps showing stem cell-related signaling pathways. Hit of the same signaling pathway identified by the EnrichR were pooled. *B*. Log p-value (PV) plotted against the pathways. * Uniqueness or significant differences between mild and severe stages. HC, hematopoietic cell (HC); SCF, stem cell factor; SP, surfactant. Other abbreviations are used in their common meanings.

### Multivariate regressions of differentially expressed exosomal proteins and clinical variables

To analyze the clinical relevance of exosomal DEPs, we analyzed the potential correlations of identified DEPs with critical clinical variables (**Figure 6**). In total, 178 proteins correlated with clinical variables (p-value < 0.05, **Supplementary file**). Among them, serine/threonine protein phosphatase 2A, fibronectin, phosphatidylethanol-amine binding protein 4, hepatoma-derived growth factor-related protein 3, and complement C_2_ showed a positive correlation with the PaO_2_/FiO_2_ ratio, i.e., the severity of ARDS. In addition, these proteins displayed a linear relationship with hemoglobin, serum HCO_3-_ (bicarbonate) level, systolic blood pressure (sysbp), and ventilation (total minute ventilation). The correlations between ARDS severity, and DEPs were analyzed and adjusted for age and gender (**Figure 6E**). The adjusted effect size of the top 22 proteins was significant (p-value < 0.05), and the overall weighted adjusted effect size (aES) favored lung regeneration (red line, p-value < 0.05). Only three downregulated proteins (P00362, P43652, and Q30134) and one upregulated protein (Q16401, S26 proteasome non-ATPase regulatory subunit 5) were shared by both mild and severe patients. Furthermore, 9 proteins were associated with ICU variables, including coagulation-free to 28-day, 60-day mortality, and 90-day mortality (adjusted p-value < 0.05). A 3-fold increase in the ratio of downregulated/upregulated proteins (9/3) was observed for severe over mild patients (3/3). Intriguingly, these 22 proteins were involved in regenerative signals (10 proteins) and growth factor pathways (5 proteins). These results demonstrate that the exosomal proteins in BAL samples could serve as predictors for the outcomes of ARDS.

**Figure 6.**
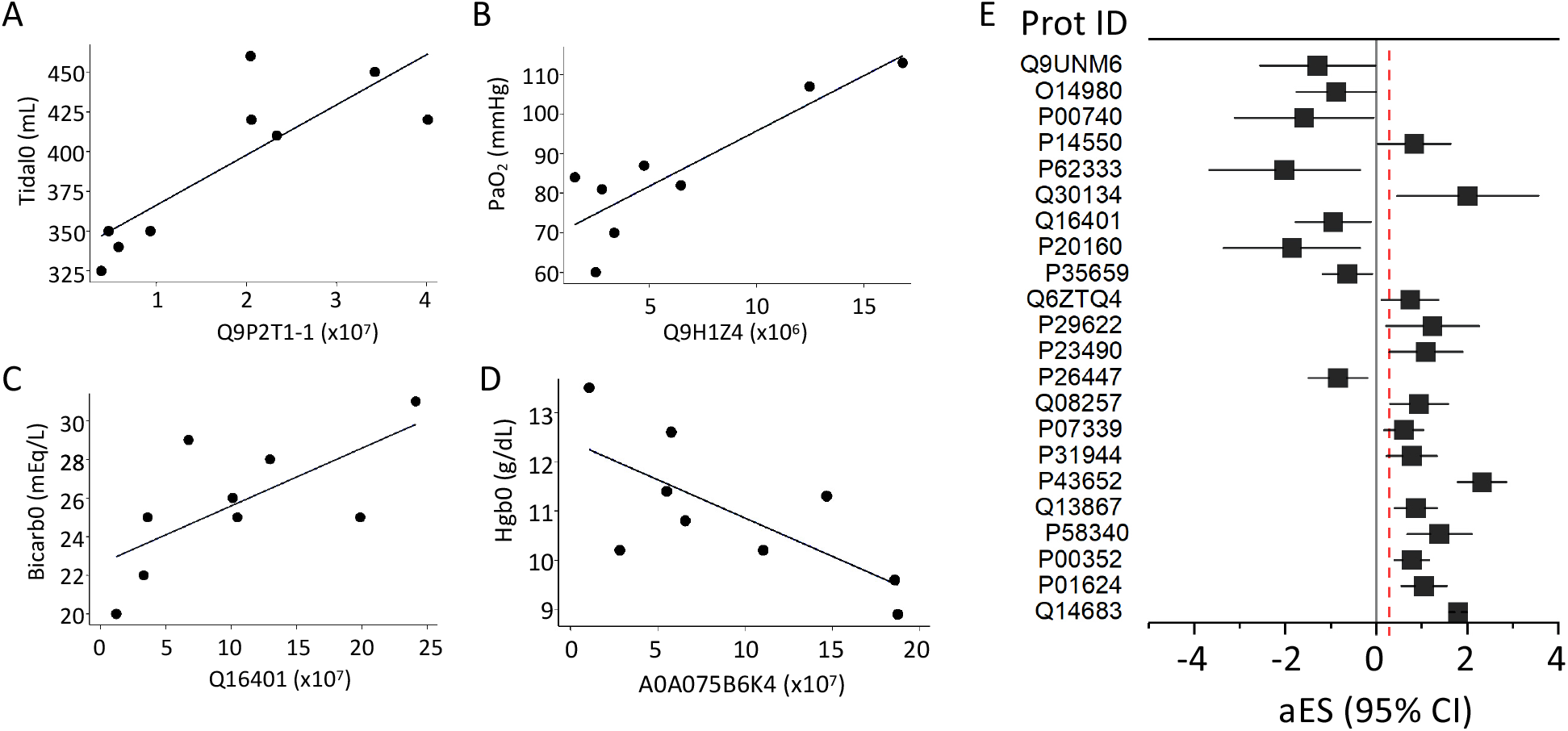
Correlations of clinical variables with differentiated BAL exosomal proteins. *A-D*. Linear regression of differentiated exosomal proteins and tidal volume (*A*), PaO_2_ (*B*), blood bicarbonate level (*C*), and hemoglobin (*D*) during their first visits. *E*. Adjusted effect size (aES) of severity (PaO_2_/FiO_2_)-associated proteins with 95% Cis (confidence intervals) adjusted for age, gender, and race. Negative aES values indicate negative associations between the protein and severity. The red line represents the overall weighted effect size positively correlating with severity.

### Divergent network between mild and severe patients

To extend our differential analysis of early and late stages of the disease, we compared the top-ten interactive networks between mild and severe groups of ARDS patients at the protein, transcriptional, and epigenetic levels (**Figure 7**). Similar to the functional annotations shown in **Figure 4**, few identical protein-protein interactions, RNA-RNA inter-regulations, and histone road maps were observed between mild and severe patients. Again, our results suggest that the niches for lung regeneration in mild patients could be significantly different from those in severe patients.

**Figure 7.**
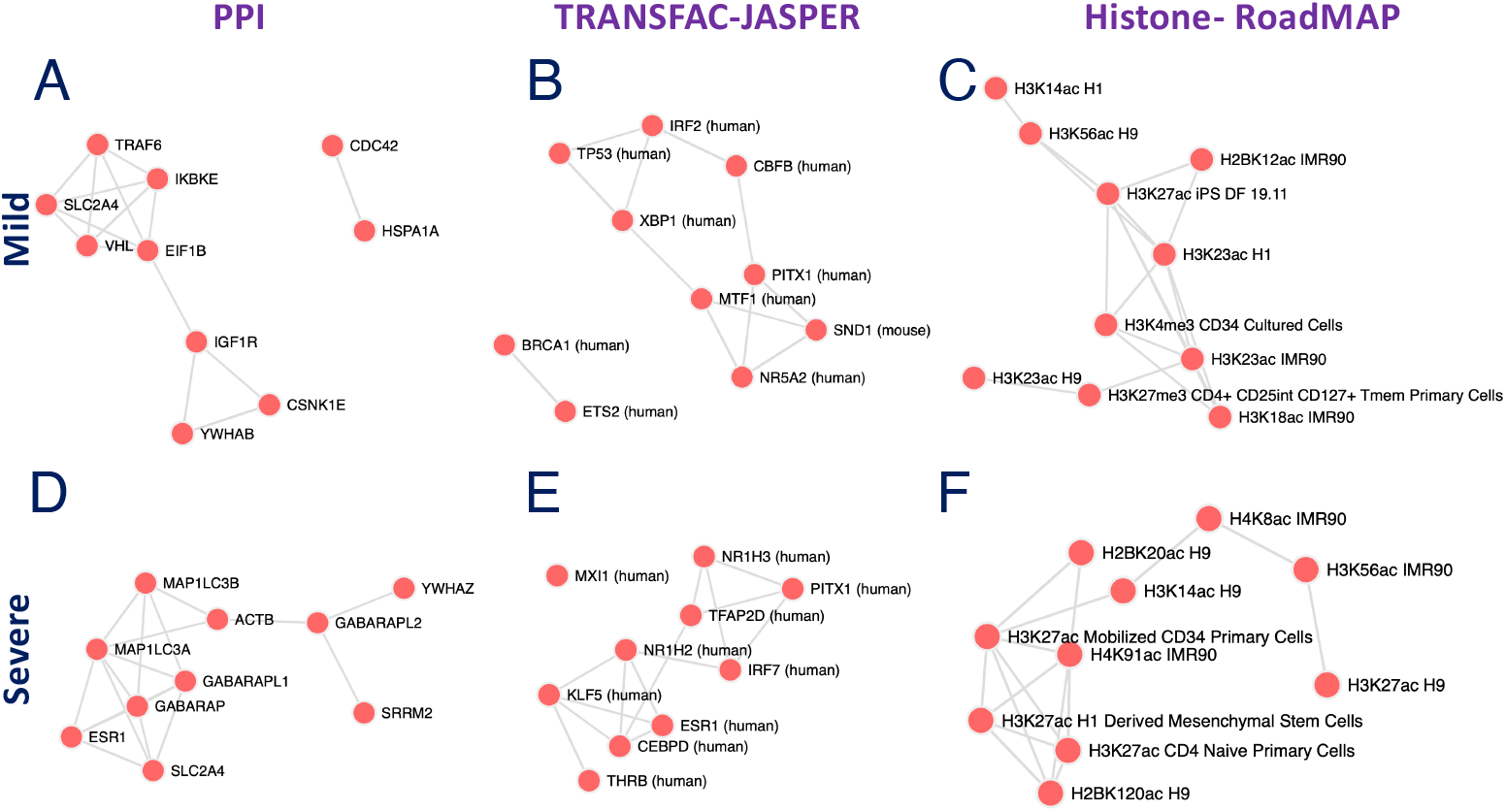
Network comparison of differentiated exosomal proteins between early and late stages of ARDS. Interactive networks were compared at the protein level (*A* & *D* by the PPI Hub Proteins), transcriptional level (*B* & *E* by the TRANSFAC-JASPER), and epigenetic level (*C* & *F* by the Epigenetics RoadMAP) between the mild and severe stages of ARDS.

### Functional enrichment for AT2 cells

AT2 cells are a major cell population that differentiates into AT1 cells to cover injured alveolar epithelium in ARDS lungs. To integrate the GO annotations, KEGG signaling pathways, and clinical diseases, we prioritized the DEPs associated with AT2 cells (**Figure 8)**. Downregulated DEPs in AT2 cells were associated with respiratory distress syndrome, chronic interstitial lung disease, and surfactant disorders. Both AT2 surfactant proteins and other AT2 biomarkers were markedly reduced. Three signaling pathways, i.e., phagosome, surfactant metabolism, and hematopoietic cell lineages, might be involved in the AT2 cell-mediated lung regeneration. In contrast, few DEPs and enrichments were observed for upregulated DEPs. These findings show that AT2 cells play a critical role in the lung regeneration by releasing exosomes and serving as a potential target to enrich regenerative niches.

**Figure 8.**
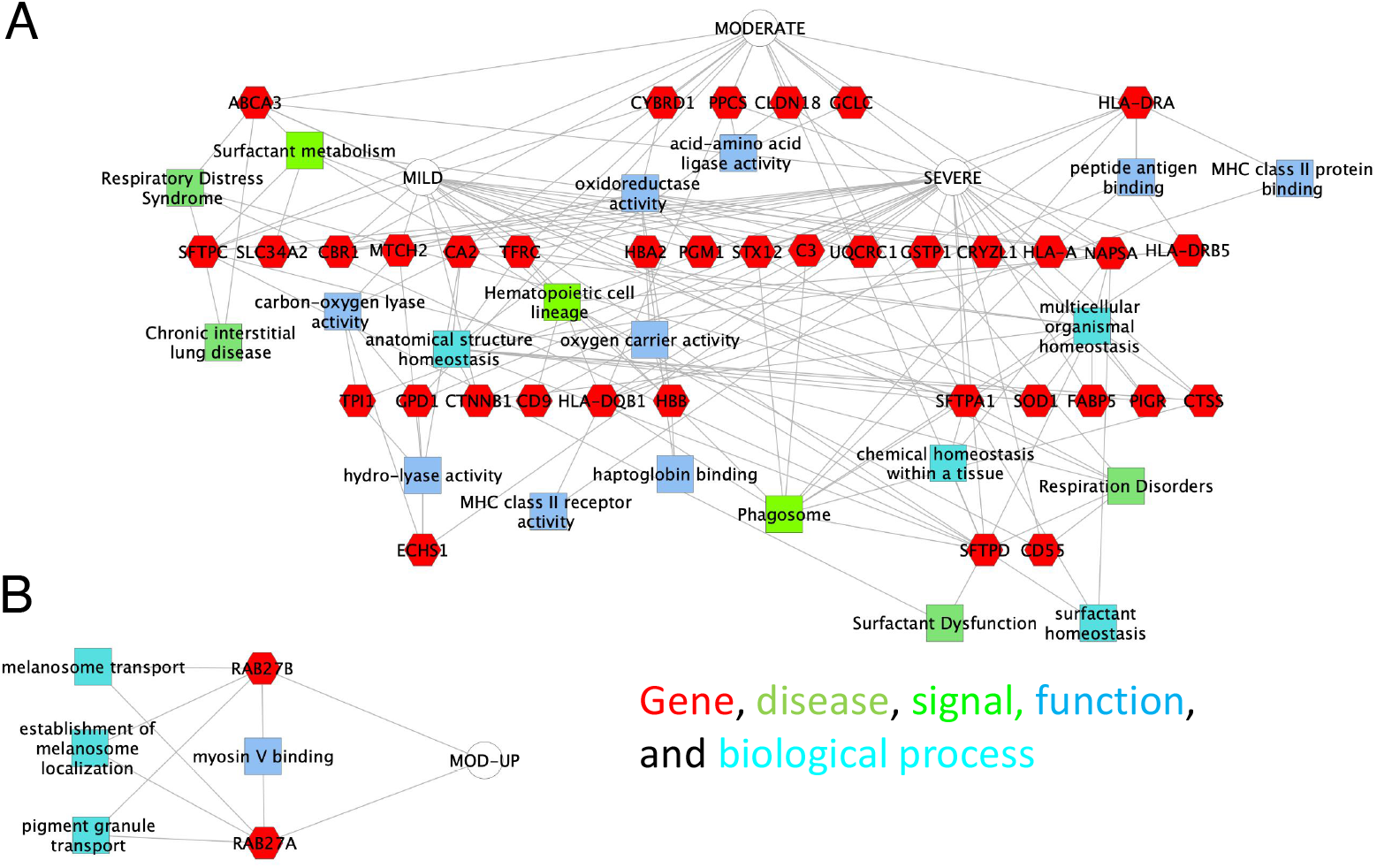
Integration of ARDS stage-specific downregulated (*A*) and upregulated (*B*) proteins, GO terms, functional annotations, and diseases in AT2 cells. The AT2 cell-prioritized ontology of differentiated exosomal proteins, including genes, diseases, signals, molecular functions, and biological processes, was analyzed using the ToppGene.

## Discussion

Considering that exosomes are crucial cargoes for the communication between lung stem cells and injured recipient cells, this study aimed to characterize the BAL exosomal proteins in ARDS patients. Our pilot study shows that more exosomal proteins are identified in the lungs of late-stage patients. Immune cells and lung epithelial stem cells are major contributors to BAL exosomes in addition to cells from other organs. Diverse functions, stem cell signals, growth factors, and immune niches were enriched between mild and severe patients. The identified DEPs were associated with key clinical variables. Severity-associated differences in protein-protein interaction, RNA crosstalk, and epigenetic network were observed between mild and severe groups of the disease. Finally, AT2 stem cells may serve as both exosome donors and recipient cells via autocrine and paracrine mechanisms.

To the best of our knowledge, this is the first study on the proteomics of BAL exosomal proteins in ARDS patients. For the first time, we identified that leukocytes (macrophages and monocytes), bone marrow stem cells (myeloid), and lung epithelial cells are three major resources of exosomal proteins in ARDS lavage. Lung regeneration, particularly, alveolar re-epithelialization could be regulated by the niches composed of exosomes released by these activated lung stem cells and immune cells. Exosomal proteins released by other organs, including the skin, gastrointestinal (GI) tract, and brain, were also identified. This is most likely a sign of their primary injury sites (trauma and GI and urinary tract infections) beyond the lung and systemic conditions (i.e., sepsis). In severe patients, exosomes released by non-lung organs could be transported to the alveoli through the blood stream via the compromised blood-gas barrier. Tracking the cell origins of BAL exosomal proteins can provide a critical alternative approach to identifying multiple organ failures and examining the barrier function. Of note, if the former scenario takes place, it cannot be detected when the barrier integrity is normal. Otherwise, if the endothelial and epithelial layers of the barrier are compromised, exosomal proteins from non-lung organs could occasionally appear in collected BAL samples.

Lung regeneration in ARDS could be regulated by the complement cascade, phagosome, neutrophil degranulation, and hemostasis. This notion is supported by the GO Biological Process of DEPs, showing that cell fate and coagulation were top-ranked. We have recently confirmed that, both in vivo and in 3D organoids, the fibrinolysis system regulates AT2-mediated re-alveolarization in injured lungs[55-57]. In ARDS lungs, fibrinolytic activity is tremendously suppressed due to elevated PAI-1 and reduced uPA and tPA[9, 58]. A downregulation of AT2-developed organoids and in vivo AT2 renewal is found in both *serpin1* transgenic and *plau* deficient preparations[55-57]. Other signaling pathways await further studies to examine their roles in lung regeneration.

The association of BAL exosomal proteins and clinical variables suggests that the identified DEPs could be potential biomarkers for the severity, outcomes, response for therapy, identification of sepsis sites, and stratification of the disease. Further identification and validation of the results in this study requires for a large cohort. Another limitation is the presence of confounding factors, including gender, race, age, and sepsis sites. A case-control-matched study will provide strong evidence for the regulation of re-alveolarization of ARDS lungs by BAL niches.

In conclusion, this study identifies novel exosomal proteins with diverse functions, signaling pathways, and cell origins in ARDS lavage samples as potential indicators of regeneration-related outcomes of the disease. Validation studies are needed to confirm the mechanisms of regenerative niches in ARDS lavage.

## Contributors

HLJ initiated the concept, HLJ, MH, RZZ, and NMX drafted the manuscript, MH, HN, SA, NVK, BS, WAT, and KPS analyzed data, all authors improved the final manuscript, and all authors approved the submission.

## Acknowledgments

This study was supported by NIH grants HL134828 (HJI) and 3RF1AG064250 (to WAT).

